# Deep mutational scanning of recent SARS-CoV-2 variants highlights changing amino acid preferences within epistatic hotspot residues

**DOI:** 10.64898/2026.03.11.711006

**Authors:** Ashley L. Taylor, Tyler N. Starr

## Abstract

Deep mutational scans across receptor-binding domains (RBDs) of diverging SARS-CoV-2 variants reveal ongoing changes to the effects of mutations, a phenomenon known as epistasis. Careful accounting for these altered mutational effects is important in viral surveillance and forecasting, and more broadly, for understanding the impacts of epistasis on real-world viral evolutionary trajectories. Using a yeast-display RBD deep mutational scanning (DMS) platform, we measure the impacts of virtually all single amino acid mutations and single-residue deletions in the Omicron KP.3.1.1 and LP.8.1 RBDs on folded RBD expression and binding affinity for the human ACE2 receptor. Our comprehensive maps reveal patterns of evolutionary accessibility and constraint at single-residue resolution and when compared to prior datasets, highlight sites whose amino acid preferences continue to change across viral variants. Notably, sites 455, 456, and 493 – which have exhibited repeated substitutions and epistatic dependencies across Omicron subvariants going back to BA.1 – again demonstrate altered patterns of mutational accessibility and constraint. Therefore, it appears that these hotspots of repeated RBD evolution have not yet converged on fixed amino acid solutions, but instead remain sites of ongoing epistatic reconfiguration. We compare our measurements of direct RBD:ACE2 affinity with recently published measurements of mutation impacts on ACE2 binding in the full quaternary spike context, which also integrates the effects of spike conformational dynamics; our analysis uncovers mutations like H505W that could favor adoption of the down/closed RBD conformation as a viral strategy for future antigenic evolution.

## INTRODUCTION

SARS-CoV-2 variants continue to emerge under evolutionary pressure to circumvent host immunity while maintaining infectivity and transmissibility [1]. These pressures drive rapid evolutionary change in the viral spike protein, which binds ACE2 receptor via the receptor-binding domain (RBD) to trigger spike conformational changes that drive fusion and cellular infection. The spike is also the target of infection- or vaccine-induced neutralizing antibodies, the most potent of which target the RBD within the same surface that it uses to engage ACE2 receptor [2,3].

Antigenic evolution of SARS-CoV-2 to escape potently neutralizing antibodies occurs via two major mechanisms. First, some mutations at sites within an antibody’s epitope reduce antibody-binding affinity and enable breakthrough infection; this mechanism is supported by the RBD’s capacity to tolerate many mutations while maintaining ACE2-binding affinity [4], minimizing the pleiotropic impacts of antibody escape [5]. Second, mutations can reduce sensitivity to neutralizing antibodies by altering spike conformational dynamics. In the quaternary spike trimer, the RBD samples two distinct conformations: a “down” (also called “closed”) conformation, which buries the RBD surface that binds ACE2 and is most susceptible to potent antibody neutralization; and an “up” (also called “open”) conformation, which is compatible with receptor-binding but exposes vulnerable epitopes to antibody inhibition [6,7]. Though bat-borne SARS-like coronaviruses are strongly biased toward RBD-down conformations [8–10], ancestral SARS-CoV-2 strains were found to adopt the RBD-up conformation with ∼50% frequency [6,7], which would enhance infection and transmissibility due to enhanced receptor engagement when pre-existing antibody pressures are absent [11,12]. Evolved SARS-CoV-2 variants have gradually biased dynamics back toward RBD-down conformations, likely as an antibody escape mechanism, though it may not be able to fully revert to the extreme RBD-down bias seen in bat-borne CoVs due to the tradeoff between antibody escape and receptor engagement that this mechanism exhibits [13,14].

Our ability to track and forecast SARS-CoV-2 evolution and understand the mechanisms described above has been aided by technological developments that expand the scale and speed of viral genomic sequencing and laboratory methods for rapid viral variant characterization [15,16]. One methodology that has emerged to fill the gap between sequencing and characterization is “deep mutational scanning” (DMS) [17], which has characterized the impacts of mutations in the SARS-CoV-2 spike and RBD on various phenotypes including ACE2-receptor binding, antibody escape, and conformational dynamics [4,18–25]. By prospectively characterizing mutational effects, DMS datasets aid in the rapid risk assessment of novel variants observed in genomic surveillance while retrospective laboratory characterization ensues. These DMS studies also inform therapeutic antibody and vaccine design, for example by identifying antibodies that are more robust to evolutionary escape [3,25,26], and can provide high-quality training or validation datasets for large model development for studies in viral and molecular evolution [20,27–30].

However, as the sequence background of the predominant SARS-CoV-2 variant changes through substitution, the impacts of mutations across RBD sites can change due to the phenomenon of epistasis [31]. We and others have documented important epistatic interactions that shape SARS-CoV-2 evolution, such as the synergistic interaction between mutations Q498R and N501Y that underlied the evolutionary success of Omicron in late 2021 [32–34], and epistasis between sites 455 and 456 that shaped the evolution of “FLip” variants in 2023 [35,36]. Epistatic perturbations are, by definition, hard to predict – therefore, there is a need to continually update the viral reference sequence on which DMS studies are performed to aid in ongoing viral surveillance and therapeutic design, and to better understand the molecular evolutionary features that shape the tempo and direction of viral variant evolution.

Two recent Omicron sub-variants that rose to global dominance were KP.3.1.1 and LP.8.1, both of which descend from Omicron BA.2.86 with shared RBD substitutions L455S (JN.1), F456L, and Q493E (**Figure 1A**). These variants began to rise in frequency midway through 2024 and maintained high levels of circulation before being replaced by related variants NB.1.8.1 and XFG midway through 2025 [37]. Notably, LP.8.1 was chosen as the vaccine strain for the northern hemisphere autumn 2025 vaccine booster. Consistent with their evolutionary success, both variants were associated with escape from serum antibody neutralization while maintaining efficient infectivity [37–39].

**Figure 1.**
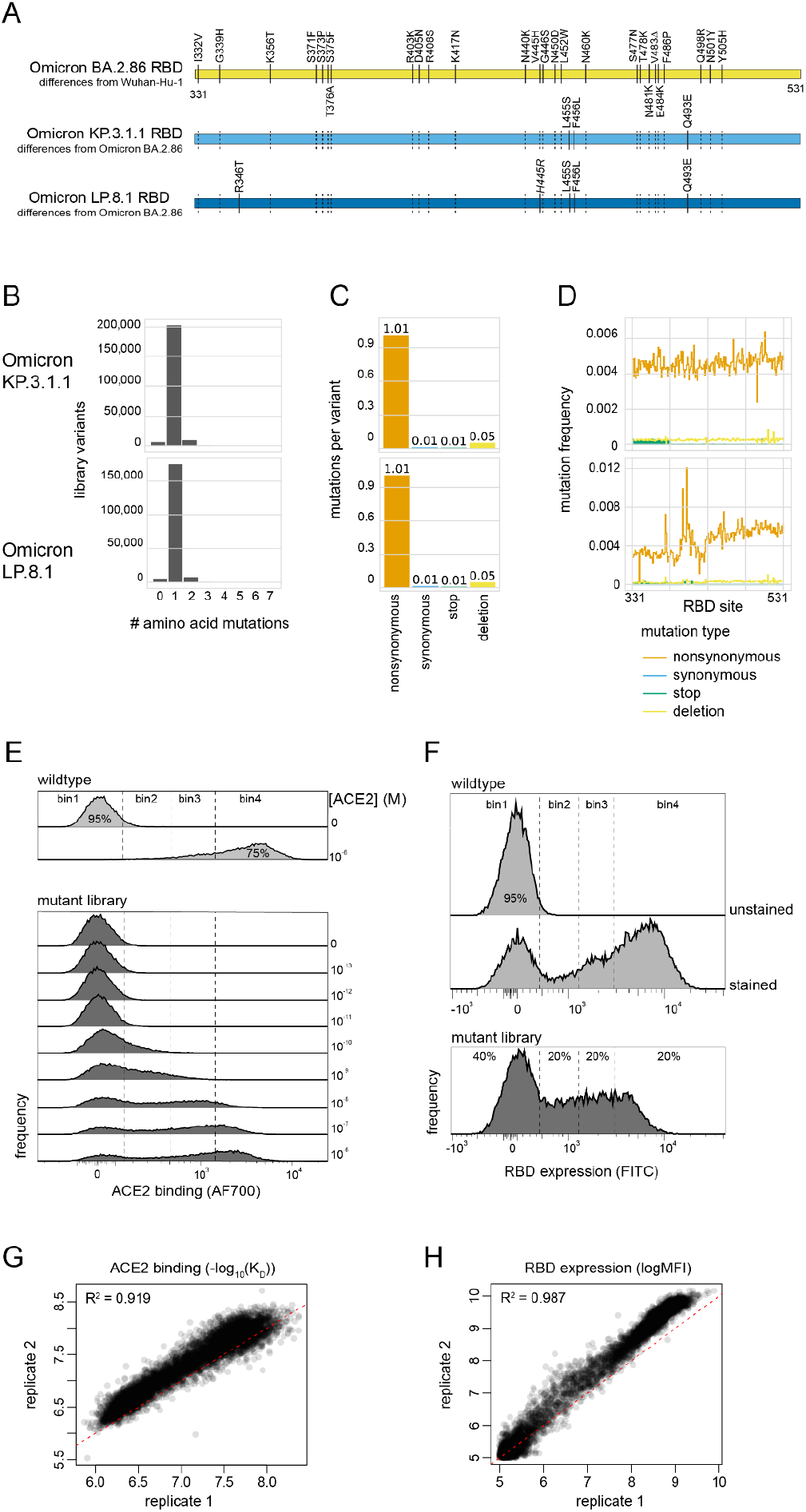
Deep mutational scanning of the SARS-CoV-2 Omicron KP.3.1.1 and LP.8.1 RBDs. (**A**) Diagram of the RBD substitutions that distinguish Omicron BA.2.86 from Wuhan-Hu-1 (top), Omicron KP.3.1.1 from BA.2.86 (middle), and Omicron LP.8.1 from BA.2.86 (bottom). Dashed lines show propagation of BA.2.86 changes to KP.3.1.1 and LP.8.1, and italicized mutation in LP.8.1 (H445R) indicates a secondary substitution at a site that previously changed from Wuhan-Hu-1 to BA.2.86. Wuhan-Hu-1 reference spike numbering is used throughout the manuscript. (**B-D**) Quality control of the KP.3.1.1 and LP.8.1 RBD site-saturation mutagenesis library as assessed by PacBio sequencing, illustrating the distribution of number of amino acid mutations per barcoded variant (B), the average number of mutations of each type across library variants (C), and the distribution of mutations across sites in the RBD over all variants (D). (**E, F**) Representative FACS gates used to sort RBD^+^ singlet cells for ACE2 binding (E) and singlet cells for RBD expression (F), which is followed by high-throughput sequencing of cells sorted into each bin. (**G, H**) Correlation in per-mutant deep mutational scanning measurements between independently barcoded replicate libraries for ACE2-binding affinity (G) and RBD expression (H) experiments. Red dash indicates the 1:1 line.

Here, we perform deep mutational scans via a yeast-surface display platform [40] to measure the impacts of all single amino acid mutations or single-codon deletions in the KP.3.1.1 and LP.8.1 RBDs on binding affinity for human ACE2 receptor. We compare our data to matched-strain DMS data for ACE2 binding in the full quaternary spike context, allowing disentanglement of mutational effects on direct RBD:ACE2 affinity and indirect RBD up/down conformation dynamics. Our data identify sites 455, 456, and 493 as ongoing hotspots of evolutionary and epistatic change, revealing newly accessible constellations of mutational accessibility at these sites that will likely continue to shape SARS-CoV-2 variant evolution.

## RESULTS

### Deep mutational scans of the KP.3.1.1 and LP.8.1 Omicron variant RBDs

We have previously described a high-throughput yeast-surface display platform for multiplexed measurement of the effects of RBD mutations on ACE2-binding affinity, which we previously used to survey the effects of mutations in SARS-CoV-2 strains Wuhan-Hu-1, Alpha, Beta, Delta, Eta, and Omicron BA.1, BA.2, BQ.1.1, XBB.1.5, EG.5, FLip, and BA.2.86 [33,36,41,42] (cites). We cloned the KP.3.1.1 and LP.8.1 RBDs (**Figure 1A**) into this yeast-display platform and constructed site-saturation mutagenesis libraries that introduce every single-amino-acid mutation or single-codon deletion at each of the 200 RBD residues. We cloned mutant libraries together with a variant-identifying N16 nucleotide barcode, and we used long-read PacBio sequencing to link RBD variants with their unique N16 barcode and confirm the integrity of our libraries. Libraries showed the intended balance of single-mutant variants (**Figure 1B**) and mutation types (**Figure 1C**), with even rates of mutation across the RBD sequence (**Figure 1D**).

We induced yeast-surface expression of our RBD variant libraries and incubated yeast-displayed libraries across a concentration gradient of monomeric human ACE2 or an antibody targeting a C-terminal epitope tag on the yeast-displayed RBD, each coupled to secondary fluorophores. We used fluorescence-activated cell sorting (FACS) to separate library variants on the basis of strength of ACE2 binding (**Figure 1E**) or RBD expression (**Figure 1F**). Cells sorted into each bin were sequenced in the N16 barcode region, enabling calculation of the impact of each mutation on ACE2-binding affinity or RBD expression based on its distribution of sequence reads across FACS bins. Deep mutational scanning measurements were conducted in duplicate from independently barcoded variant libraries, demonstrating strong correlation in the measured effects of mutations (**Figure 1G, H**).

Heatmaps illustrating the impact of each RBD mutation on ACE2-binding affinity and RBD expression, which are very similar between the closely related KP.3.1.1 and LP.8.1 variants, are presented in **Figure 2**. An interactive version of these heatmaps, integrated with measurements in prior SARS-CoV-2 variant RBDs, is available at: https://tstarrlab.github.io/SARS-CoV-2-RBD_DMS_Omicron-KP3-LP8/RBD-heatmaps/.

**Figure 2.**
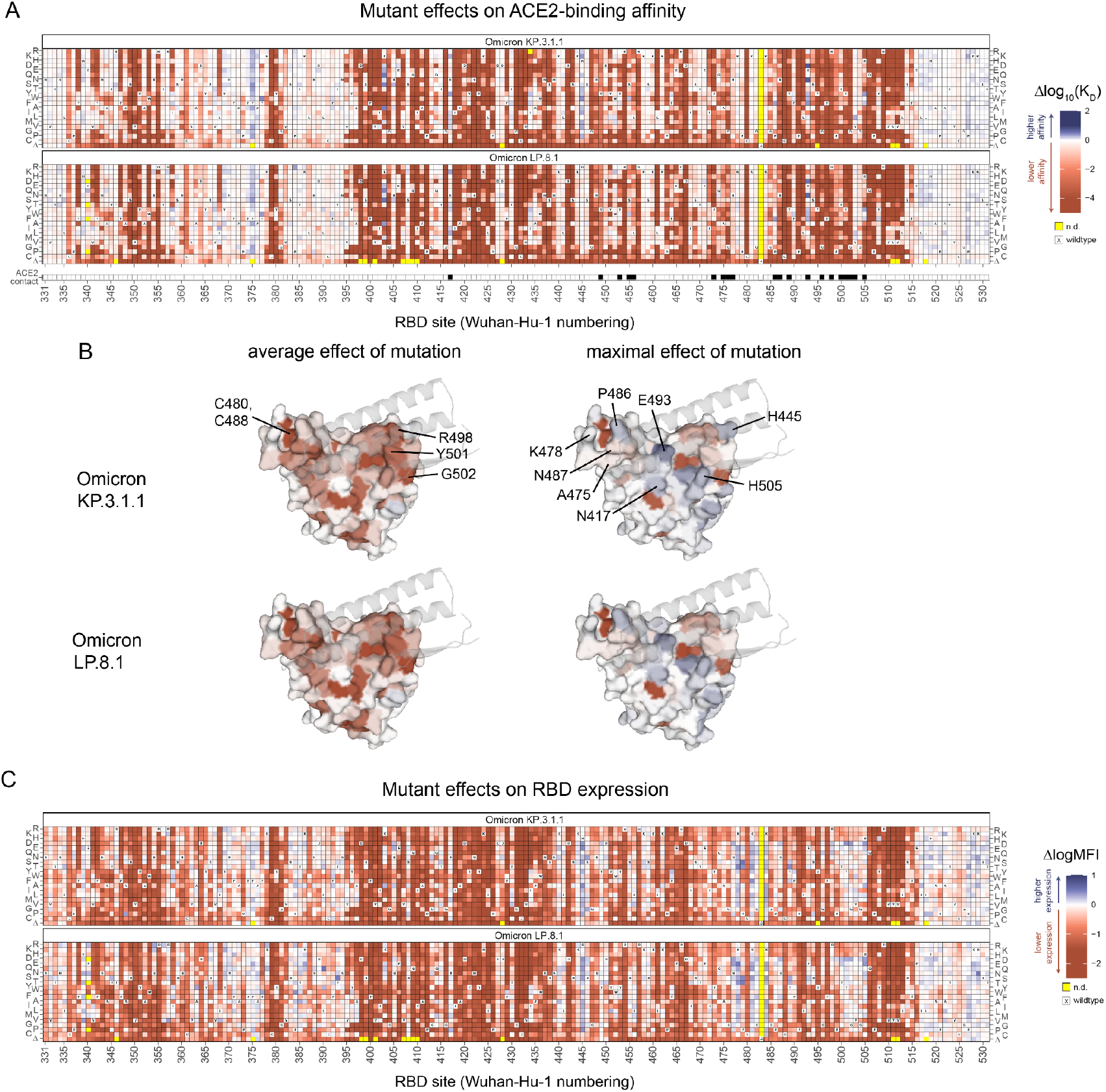
Effects of mutations in the KP.3.1.1 and LP.8.1 receptor-binding domain on ACE2 binding and RBD expression. (**A**) Heatmap illustrating the impacts of all mutations in the KP.3.1.1 and LP.8.1 RBD on ACE2-binding affinity as determined from FACS-seq experiments with yeast-displayed RBD mutant libraries. ACE2 contact residues (black squares, bottom) defined as RBD residues with non-hydrogen atoms <5Å from ACE2 in the BA.2.86 RBD structure (PDB 8QSQ [46]). Raw data available in Supplemental Data 1. (**B**) Deep mutational scanning data from (A) mapped to the ACE2-bound BA.2.86 RBD structure (PDB 8QSQ [46]), illustrating the average effect of mutations at a site (left), and the maximal effect of any mutation at a site (right). Sites of interest are labeled. ACE2 (key motifs only) is shown as transparent gray cartoon. (**C**) Heatmap illustrating the impacts of all mutations in the KP.3.1.1 and LP.8.1 RBD on yeast-surface expression levels, a proxy for folding and expression efficiency. Raw data available in Supplemental Data 1. An interactive version of these heatmaps alongside previously assayed SARS-CoV-2 variant RBDs is available at https://tstarrlab.github.io/SARS-CoV-2-RBD_DMS_Omicron-KP3-LP8/RBD-heatmaps/.

Our maps show expected patterns of heterogeneity in tolerance to mutation versus constraint across RBD positions. For example, all mutations to cysteines involved in disulfide bridges, such as those at positions 480 and 488, are deleterious. We also see strong constraint among some ACE2-contact residues. For example, mutations are not well tolerated at residue G502, where steric hindrance with ACE2 prevents the introduction of any side chain [4]. Mutations are also poorly tolerated at residues R498 and Y501; these two residues exhibited synergistic epistasis in the emergence of Omicron [32–34], and have been subsequently “entrenched” [43,44] by co-occurring substitutions in Omicron that are dependent on the substituted R498 and/or Y501 residues, which may hinder subsequent evolutionary change at these sites.

Despite the presence of many deleterious mutations, various amino acid changes are tolerated or can even enhance affinity for human ACE2 receptor, including at sites 417, 455, 456, 486, and 505 whose substitutions have defined success of prior variants of concern. Of note, residues 475, 478, and 487 – which were recently highlighted as sites of strong escape from JN.1-like antibody responses [45] – can tolerate many amino acid changes without strong pleiotropic constraint for ACE2 binding, which may support the ongoing evolution of variants such as the currently prevalent XFG (which carries N487D) and NB.1.8.1 (which carries K478I).

### Disentangling the role of ACE2 affinity and conformational dynamics on receptor binding

In addition to mutations directly impacting binding affinity between RBD and ACE2, mutations can also indirectly modulate ACE2 binding in the quaternary spike context by altering the propensity with which RBDs adopt the “up” conformation (competent for receptor-engagement but vulnerable to neutralizing antibody binding) versus pack “down” (where RBD cannot bind receptor but also hides critical epitopes from host antibodies). Mutations that bias RBD toward the down conformation can escape a wider swath of antibodies than mutations within individual antibody epitopes and are thus important to survey.

Previous work from Dadonaite et al. employed a pseudovirus deep mutational scanning platform to measure the impacts of mutations in the full spike trimer [45], in contrast to isolated RBD studied here. In the whole-spike context, one can perform neutralization assays with soluble ACE2 protein as a proxy for ACE2-binding strength [23,47,48], which integrates direct RBD:ACE2 affinity and RBD up/down conformational dynamics. Dadonaite et al. identified that many mutations distal to the RBD:ACE2 interface can impact ACE2 binding via indirect conformational effects, evidenced by the anticorrelated effects of these mutations on potency of ACE2 binding and serum neutralization. However, this method cannot easily identify the conformational impacts of mutations to RBD sites close to the RBD:ACE2 interface, because an observable effect on ACE2 binding for these mutations could come from direct effects on affinity, indirect effects on conformation, or some combination of the two. We reasoned that comparing whole-spike pseudoviral DMS with our current RBD DMS could cast further light on the effects of mutations within the ACE2-binding interface on spike conformational dynamics.

To disentangle these mechanisms, we examined the correlation between mutant effects in KP.3.1.1 on ACE2 binding from whole-spike DMS (incorporating both direct-affinity and indirect-conformation effects) and RBD DMS (incorporating just direct-affinity) (**Figure 3A**). As expected, for sites distal from the RBD:ACE2 interface (>8Å), virtually none of the mutant impacts on ACE2 binding in spike trimer are explained by direct-affinity effects (R^2^ = 0.001), indicating that these mutations impact ACE2 binding solely through indirect effects (i.e., conformational dynamics). In contrast, for sites that are proximal to (5-8Å) or directly at (<5Å) the ACE2 interface, the impacts of mutations on direct-affinity explain the majority of variation in mutant effects on ACE2 engagement in the spike trimer (R^2^ = 0.653 and 0.848 for proximal and direct-contact residues, respectively), indicating weak (contact) or modest (proximal) contributions from conformational effects.

**Figure 3.**
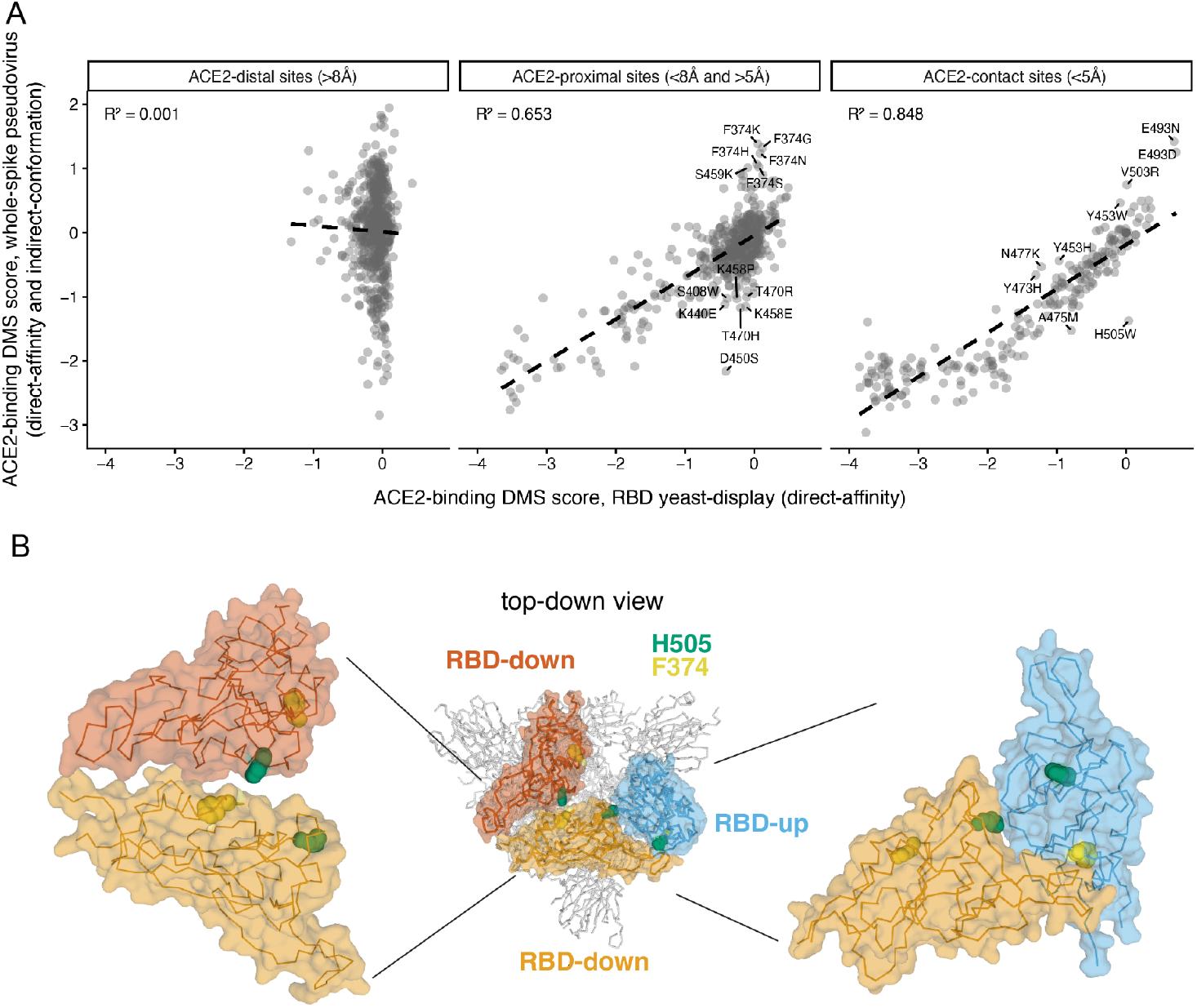
Disentangling mutational effects on direct ACE2-binding affinity and conformational dynamics. (**A**) Correlation between KP.3.1.1 mutant effects on ACE2 binding in the pseudovirus whole-spike DMS study of Dadonaite et al. (change in potency of pseudoviral neutralization via soluble ACE2 protein) [45] versus the yeast-display RBD DMS study presented here (change in binding affinity from FACS-seq titration). Correlations are separated based on residue distance from the ACE2 interface in the ACE2-bound BA.2.86 RBD structure (PDB 8QSQ [46]). Mutations whose effect on spike-mediated pseudoviral entry was < -2 units per the assay of Dadonaite et al. were not included because it is difficult to reliably measure soluble-ACE2-inhibition of entry when entry is poor. (**B**) Structural context of residues F374 and H505 in the spike trimer (PDB 9ELH [52]). Center, overall top-down view of the spike trimer with two RBDs in the “down” and one in the “up” conformation. Left, zoom-in on the down-down interface of RBD packing; right, zoom-in on the up-down interface of RBD packing.

Despite direct-affinity effects explaining the majority of variation in whole-spike ACE2 binding among ACE2-proximal and -contact residues, our analysis highlights outlier mutations that may impact RBD up/down dynamics above and beyond direct-affinity effects. For example, mutations to ACE2-proximal residue F374 do not have a large impact on direct ACE2-binding affinity, but several mutations enhance ACE2 binding in spike trimer suggesting these mutations bias RBDs toward the up conformation. Residue 374 packs with the α-helix spanning residues ∼365-370, whose glycosylation state and conformation has previously been shown to gate the up/down conformational transition [49,50]. In fact, this α-helix was the site of the substitution unique to SARS-CoV-2 that knocks out the N370 glycan conserved among all other SARS-like coronaviruses, underlying ancestral SARS-CoV-2’s infectivity by promoting RBD-up conformations [11,12]. This surface was also the site of substitutions S371L/F, S373P, and S375F that enhanced immune escape in the emergence of Omicron BA.1 and BA.2 through putative modulation of up/down dynamics [51].

Among direct ACE2-contact residues, H505W stands out as a mutation that decreases receptor binding in the spike DMS dataset despite having no meaningful effect on ACE2-binding affinity in the yeast-display data; this pattern suggests that H505W biases RBDs toward the down conformation, consistent with its reduction of neutralization by serum antibodies [45]. Residue 505 creates packing contacts at RBD:RBD interfaces in both up and down RBD conformations (**Figure 3B)**, and some mutations to this site also influence direct ACE2-binding affinity (**Figure 2**), highlighting the complex interplay mutations at site 505 may have on receptor binding. Mutations to E493 also stand out as outliers that may contribute to ACE2 binding by synergistic effects on RBD conformation and direct affinity; however the propensity of mutations at residue E493 to fall off of the correlation line at its upper limits could also reflect a simple difference in scaling of dynamic ranges between these two DMS assays.

### Epistatic shifts in mutational effects

We next examined the extent to which the effects of mutations as measured in KP.3.1.1 and LP.8.1 differ from those measured in their BA.2.86 ancestor due to epistasis. We computed a sitewise metric of divergence between the vectors of affinities measured for the 20 amino acids or a deletion between any pair of RBD backgrounds (**Figure 4**; interactive visualization of all variant-pairs and sites available at https://tstarrlab.github.io/SARS-CoV-2-RBD_DMS_Omicron-KP3-LP8/epistatic-shifts/). This “epistatic shift” metric scales from 0 to 1, with 0 indicating a site where the measured affinities of each amino acid mutant are identical between RBD variants and 1 indicating a site where the distributions are entirely dissimilar [33].

**Figure 4.**
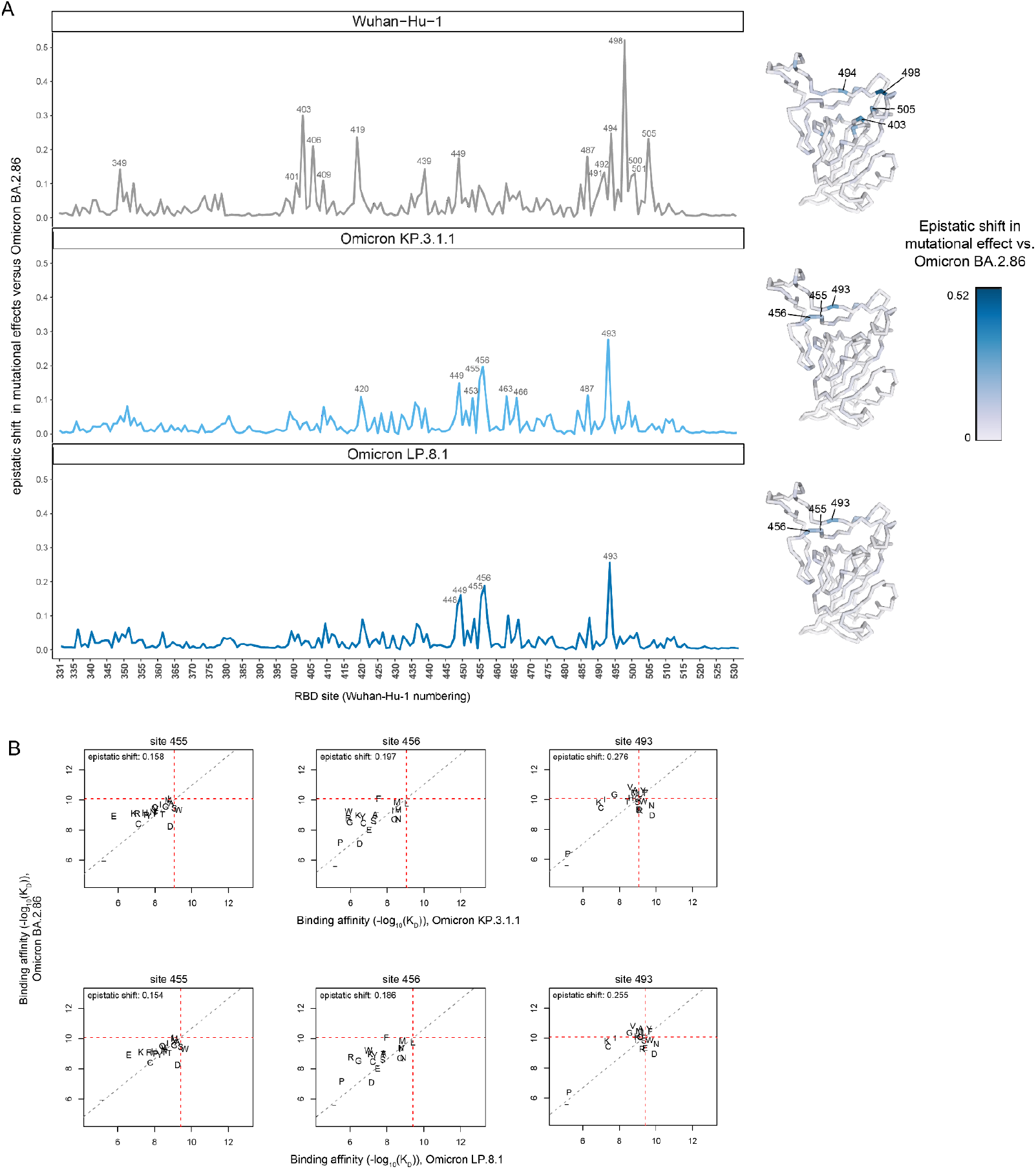
Epistatic shifts in mutational effects on ACE2 binding. (**A**) Epistatic shift in the effects of mutations on ACE2 binding at each RBD position as measured in the Wuhan-Hu-1 (previously reported in [33]), KP.3.1.1 or LP.8.1 background compared to those previously measured in Omicron BA.2.86 (previously reported in [41]). (**B**) Mutation-level plots of epistatic shifts between BA.2.86 and KP.3.1.1 or LP.8.1 at sites of interest. Each scatterplot shows the measured ACE2-binding affinity of each amino acid (plotting character, – indicates deletion character) in the BA.2.86 versus KP.3.1.1 or LP.8.1 backgrounds. Red dashed lines mark the respective wildtype RBD affinities on each axis, and the gray dashed line indicates the additive (non-epistatic) expectation. Interactive plots enabling the comparison of all SARS-CoV-2 variants and scatterplots for all RBD sites are available at https://tstarrlab.github.io/SARS-CoV-2-RBD_DMS_Omicron-KP3-LP8/epistatic-shifts/.

We previously observed widespread epistatic shifts in the effects of mutations at many sites between the ancestral strain Wuhan-Hu-1 and strains like Alpha or Omicron that carry N501Y, a strong epistatic modifier substitution (**Figure 4A**, top) [33,41]. Though not as strong or widespread as these prior epistatic shifts, we observe ongoing epistatic shifts in the effects of mutations at select sites in the evolution from BA.2.86 to KP.3.1.1 and LP.8.1 (**Figure 4A**, middle and bottom). The strongest site of perturbation is site 493, where further examination of individual mutational effects shows many sign-epistatic changes (**Figure 4B**). For example, mutation to V493 is affinity-enhancing in BA.2.86 but deleterious in KP.3.1.1 and LP.8.1, whereas mutation to D493 shows the opposite pattern. These patterns reflect biochemical entrenchment [43,44], wherein accommodation of E493 in the evolution of KP.3.1.1 and LP.8.1 leads to correlated accommodation of biochemically similar amino acids D (which shares the negatively charged carboxylic acid moiety with glutamate) and N (which is sterically similar to glutamate, with a polar amide group in place of the carboxylic acid moiety). Although they don’t show as widespread of sign-epistatic changes between BA.2.86 and KP.3.1.1 or LP.8.1, sites 455 and 456 also show strong epistatic shifts during recent SARS-CoV-2 evolution.

### Dynamic epistatic shifts over SARS-CoV-2 variant evolution

Sites 455, 456, and 493 have attracted attention due to their repeated substitution during Omicron subvariant evolution over the past two years (**Figure 5A**), potentiated by epistatic interactions between mutations at these sites [35,36,42]. For example, we previously showed that the Q493E mutation was deleterious for ACE2 binding for all SARS-CoV-2 variants until it emerged in KP.3 coincidental with the L455S and F456L substitutions, which reverse the sign of the effect of Q493E on ACE2-binding affinity [42].

**Figure 5.**
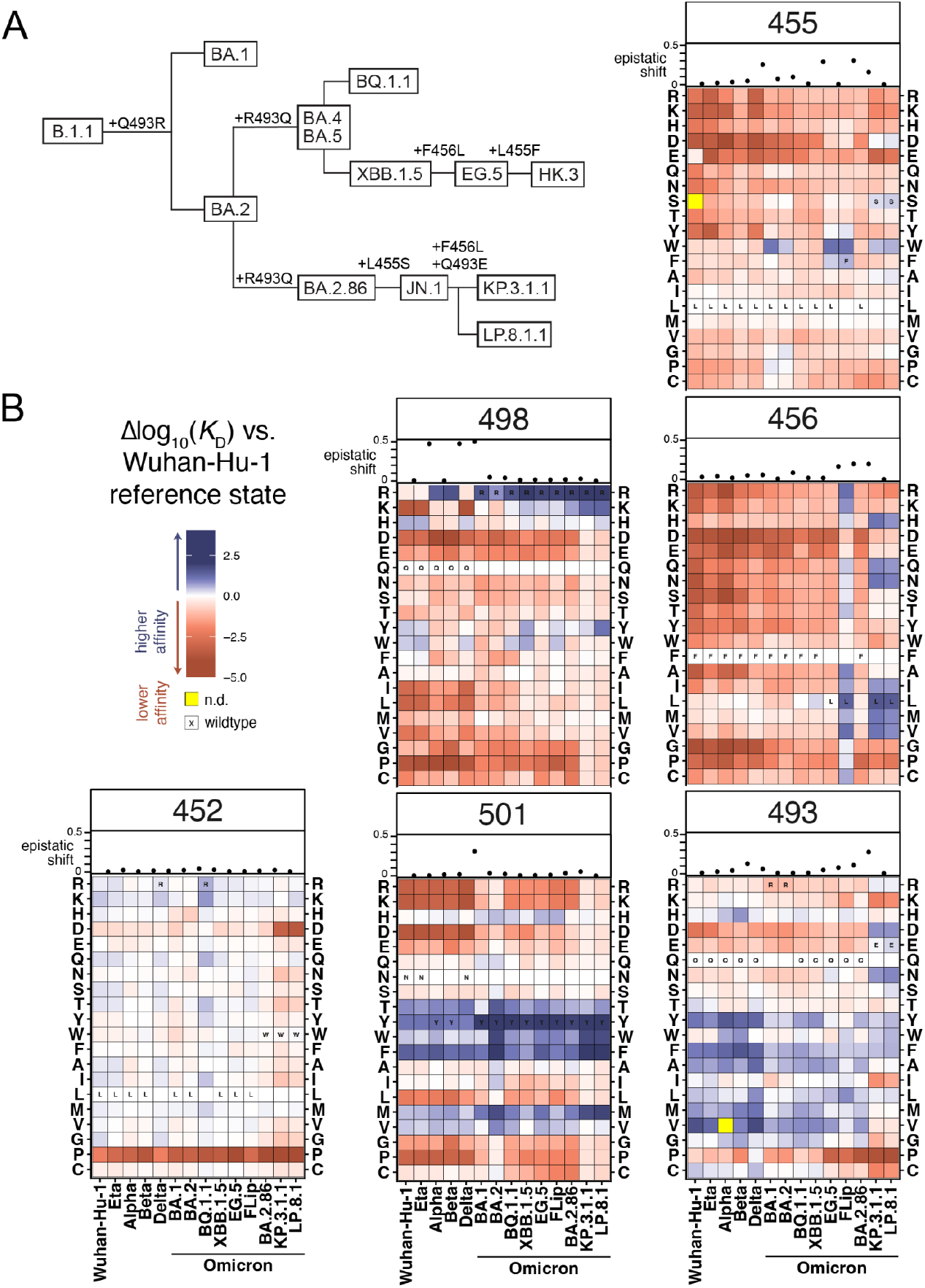
(**A**) Cladogram showing relationships among select SARS-CoV-2 Omicron variants, with amino acid substitutions at positions 455, 456, and 493 indicated (other mutations not shown). (**B**) Heatmaps of mutational effects at select sites of interest across assayed SARS-CoV-2 RBD variants. For each heatmap, mutational effects (Δlog_10_*K*_D_) are polarized to be expressed relative to the same reference amino acid state (the ancestral Wuhan-Hu-1 identity) in each column so preferences can be visualized side-by-side without changes in wildtype identity altering the reference state. Plots above each heatmap show the summary epistatic shift metric for that site between each pair of backgrounds from left to right.

To better visualize changes in evolvability and constraint at this trio of positions over time, we created composite heatmaps that illustrate the effects of mutations at a site across all SARS-CoV-2 RBD backgrounds in which we have completed DMS measurements (**Figure 5B**). For these heatmaps, we re-polarize all mutations to be expressed relative to the ancestral Wuhan-Hu-1 amino acid reference state at that position, enabling visual comparison of sitewise mutational profiles when wildtype residues differ between variant backgrounds. Above each heatmap, we show the epistatic shift between each pair of backgrounds from left to right, which more or less reflects the temporal order of variant emergence over the course of the pandemic. For comparison, we also show composite heatmaps at residues 452, 498, and 501—sites that have shown similar tempos of substitution over SARS-CoV-2 variant evolution as sites 455, 456, and 493.

Despite repeatedly substituting across SARS-CoV-2 variants, amino acid preferences at site 452 have been quite consistent across RBD backgrounds. Substitutions at this site may therefore represent changing selective pressures on the site from e.g. antigenic targeting or spike conformational dynamics. On the other hand, epistatic shifts at sites 498 and 501 between variants were quite strong during early SARS-CoV-2 variant evolution. However, since the fixation of the synergistic pair of R498 and Y501 in Omicron BA.1, amino acid preferences at these positions have remained relatively stable—though it is worth noting the presence of mutations like R498Y, Y501F, or Y501M that maintain strong ACE2-binding affinity and could unlock future epistatic evolutionary trajectories at the complementary site.

In contrast to sites 498 and 501 which showed strong but acute epistatic shifts, residues 455, 456, and 493 have exhibited enduring epistatic shifts over the course of Omicron evolution that continue to yield novel amino acid preferences. For example, mutations at residue 456 to amino acids H, Q, N, I, M, and V are newly enabled in recent Omicron sub-variants, as are substitutions like E493D and E493N. The rapidly and consistently fluctuating amino acid preferences at these positions make it challenging to predict the direction of future evolution at these positions, which is important given their importance for potential for escape from clinically relevant monoclonal antibodies such as SA55 and VIR-7229 [26,53].

## DISCUSSION

The evolutionary forces driving spike evolution have changed over the course of the pandemic [1]: for the first two years following SARS-CoV-2 emergence, selection drove substitutions like D614G and N501Y that enhanced transmission by altering spike affinity for receptor, stability, or fusogenicity. Following the initial spread of SARS-CoV-2 and induction of widespread population immunity, selection has shifted to favor substitutions that reduce antibody-mediated neutralization with the ongoing constraint to maintain receptor engagement and fusion. The large number of mutations tolerated in the spike RBD while maintaining high-affinity binding to ACE2, as revealed by deep mutational scans, is likely to contribute to SARS-CoV-2’s capacity for antigenic evolution by minimizing deleterious side-effects of key immune escape mutations.

One limitation of our approach is the focus on single-mutant effects, which can create challenges for predicting the combined effects of multiple mutations that typically accrue in newly emerging variants. Unfortunately this remains an ongoing challenge due to the vast space of possible combinatorial mutants and relative sparsity of consequential epistasis within these combinatorial landscapes [54,55]. Nonetheless, our single-mutant scans across variant backgrounds can be leveraged for modeling of combinatorial effects [20], and our identification of “hotspots” of epistatic interaction opens up the potential for high-yield combinatorial explorations. Another limitation is our focus on an isolated-RBD platform for querying RBD:ACE2 interaction. Fortunately, complementary deep mutational scanning platforms exist to query mutational impacts in the full quaternary spike context [23,45]; as shown here, combining the biological complexity of these platforms with the precision of our platform can enable new mechanistic insights, for example improving our ability to disentangle mutational effects on direct ACE2 binding and indirect effects on RBD conformational dynamics.

As the evolutionary forces driving spike change have changed, so have the fundamental effects of the mutations on which selection is acting due to epistatic shifts in the impacts of mutations such as those documented here. By conducting deep mutational scans across major SARS-CoV-2 variants from Wuhan-Hu-1 to present, we have identified key epistatic interactions that defined variant success through different dynamics. For example, epistasis between positions 498 and 501 dominated early SARS-CoV-2 evolution through the initial emergence of Omicron [33]. In contrast, we have found that positions 455, 456, and 493 exhibit more persistent and recurrent epistatic shifts that continue to shape the evolution of new Omicron sub-variants. Our newest deep mutational scans in Omicron KP.3.1 and LP.8.1 reveal yet new patterns of amino acid preferences at this trio of residues; it therefore seems as though evolution might continue to sample new combinations of mutations at these positions for more time to come.

## MATERIALS & METHODS

### Mutant libraries

We cloned yeast codon-optimized RBD sequences (amino acids N331 – T531 by Wuhan-Hu-1 reference numbering, the numbering index we use throughout the manuscript) from Omicron KP.3.1.1 and LP.8.1 into a yeast surface display plasmid as described [41]. Parental plasmid and associated sequence maps are available from Addgene (Addgene ID in process) and https://github.com/tstarrlab/SARS-CoV-2-RBD_DMS_Omicron-KP3-LP8/blob/main/data/336_pETcon_SARS2_KP.3.gb, https://github.com/tstarrlab/SARS-CoV-2-RBD_DMS_Omicron-KP3-LP8/blob/main/data/420_pETcon_SARS2_LP.8.1.gb.

Site-saturation mutagenesis libraries spanning all 200 positions in the KP.3.1.1 and LP.8.1 RBDs were produced by Twist Bioscience. We programmed the introduction of precise codon mutations to encode the 19 possible amino acid mutations at each RBD position and a single-codon deletion. To ensure an adequate level of relevant control variants in the library, stop codon mutations were programmed to be introduced at every other position for the first 50 positions, and wildtype codons were specified at every other position for the first 100 positions. Libraries were delivered as dsDNA oligonucleotides with constant flanking sequences. The “mutant RBD fragment” sequence delivered for KP.3.1.1 as an example (where uppercase letters denote mutated region) is:

~~~
tctgcaggctagtggtggaggaggctctggtggaggcggccgcggaggcggagggtcggctagccatatgAACGTTACCAACTTGTGTCCAT
TCCATGAAGTTTTCAATGCTACTAGATTCGCTTCTGTTTACGCTTGGAATAGAACTAGAATCTCTAACTGCGTTGCTGACTATTCTGTCTTG
TACAATTTTGCTCCATTCTTCGCTTTCAAGTGCTATGGTGTTTCTCCAACTAAGTTGAACGATTTGTGTTTCACCAACGTTTACGCCGATTC
CTTTGTTATTAAAGGTAACGAAGTCTCCCAAATTGCTCCAGGTCAAACTGGTAATATTGCCGATTACAATTACAAGTTGCCAGATGATTTCA
CCGGTTGTGTTATTGCTTGGAACTCTAACAAGTTGGATTCTAAGCATTCTGGCAACTACGATTACTGGTACAGGTCTTTGCGTAAGTCCAAA
TTGAAGCCATTCGAAAGAGATATTTCCACCGAAATCTATCAAGCTGGTAACAAGCCATGTAAAGGTAAAGGTCCAAACTGTTACTTCCCATT
GGAATCTTACGGTTTCAGACCAACTTATGGTGTTGGTCATCAACCATACAGAGTTGTTGTTTTGTCTTTCGAGTTGTTGCATGCTCCAGCTA
CTGTTTGTGGTCCAAAGAAATCTACTctcgaggggggcggttccgaacaaaagcttatttctgaagaggacttgtaatagagatctgataac
aacagtgtagatgtaacaaaatcgactttgttcccactgtacttttagctcg
~~~

A second dsDNA fragment encoding constant flanks and a randomized N16 barcode was produced via PCR off of the parental vector with primer-based sequence additions (primers described in [4,56]). This “barcode fragment” sequence is:

~~~
cgactttgttcccactgtacttttagctcgtacaaaatacaatatacttttcatttctccgtaaacaacatgttttcccatgtaatatcctt
ttctatttttcgttccgttaccaactttacacatactttatatagctattcacttctatacactaaaaaactaagacaattttaattttgct
gcctgccatatttcaatttgttataaattcctataatttatcctattagtagctaaaaaaagatgaatgtgaatcgaatcctaagagaatta
atgatacggcgaccaccgagatctacactctttccctacacgacgctcttccgatctNNNNNNNNNNNNNNNNgcggccgcgagctccaatt
cgccctatagtgagtcgtattacaattcactgg
~~~

The mutant RBD fragment and barcode fragment were combined with NotI/SacI-digested parental plasmid backbone via HiFi Assembly. An example of the structure of the final assembled library (in the KP.3.1.1 background) is available on GitHub: https://github.com/tstarrlab/SARS-CoV-2-RBD_DMS_Omicron-KP3-LP8/blob/main/data/336lib_pETcon_SARS2_KP.3.gb. Assembled library plasmids were electroporated into *E. coli* (NEB 10-beta, New England Biolabs C3020K), and plated at limiting dilutions on LB+ampicillin plates. For each library, duplicate plates corresponding to an estimated bottleneck of ∼80,000 cfu for each library were scraped and plasmid purified, such that each of the 4000 RBD mutations are linked to an average of 20 barcodes. For positions that failed mutagenesis QC from Twist, in-house mutagenesis pools for each position were constructed via PCR with NNS mutagenic primers, Gibson assembled, plated to approximately ∼400 cfu per position, and plasmid purified and pooled with the primary plasmid library. Plasmid libraries are available from Addgene (Addgene ID in process). Plasmid libraries were transformed into the AWY101 yeast strain [57] at 10-μg scale according to the protocol of Gietz and Schiestl [58], and aliquots of 9 OD*mL of yeast outgrowth were flash frozen and stored at -80°C.

As described previously [4,33,41,56], we sequenced NotI-digested plasmid libraries on a PacBio Revio to generate long sequence reads spanning the N16 barcode and mutant RBD coding sequence. The resulting circular consensus sequence (CCS) reads are available on the NCBI Sequence Read Archive (SRA), BioProject PRJNA770094, BioSample SAMN56444854. PacBio CCSs were processed using alignparse version 0.6.0 [59] to call N16 barcode sequence and RBD variant genotype and filter for high-quality sequences. Analysis of the PacBio sequencing indicates that all but 2 of the intended 4000 RBD mutations were sampled on ≥1 barcode in the KP.3.1.1 libraries, and all but 11 were sampled on ≥1 barcode in the LP.8.1 libraries with even coverage (Figure 1D). Complete computational pipelines and summary plots for PacBio data processing and library analysis are available on GitHub: https://github.com/tstarrlab/SARS-CoV-2-RBD_DMS_Omicron-KP3-LP8/blob/main/results/summary/process_ccs_KP3.md and https://github.com/tstarrlab/SARS-CoV-2-RBD_DMS_Omicron-KP3-LP8/blob/main/results/summary/process_ccs_LP8.md. Final barcode-variant lookup table is available on GitHub: https://github.com/tstarrlab/SARS-CoV-2-RBD_DMS_Omicron-KP3-LP8/tree/main/results/variants.

### Deep mutational scanning for ACE2-binding affinity

The effects of mutations on ACE2 binding affinity were determined via FACS-seq titration assays [4,33,36,40,41]. Titrations were performed in duplicate on independently barcoded mutant libraries. Frozen yeast libraries were thawed, grown overnight at 30°C in SD -Ura -Trp media (8 g/L Yeast Nitrogen Base, 2 g/L -Ura -Trp Synthetic Complete dropout powder and 2% w/v dextrose), and backdiluted to 0.67 OD600 in SG -Ura -Trp + 0.1%D (recipe as above but with 2% galactose and 0.1% dextrose in place of the 2% dextrose) to induce RBD expression, which proceeded for 22-24 hours at 19°C with gentle mixing.

Induced cells were washed with PBS-BSA (BSA 0.2 mg/L), split into 16-OD*mL aliquots, and incubated with biotinylated monomeric human ACE2 protein (ACROBiosystems AC2-H82E8) across a concentration range from 10^-6^ to 10^-13^ M at 1-log intervals, plus a 0 M sample. Incubations equilibrated overnight at room temperature with gentle mixing. Yeast were washed twice with ice-cold PBS-BSA and fluorescently labeled for 1 hr at 4°C with 1:100 FITC-conjugated chicken anti-Myc (Immunology Consultants CMYC-45F) to detect yeast-displayed RBD protein and 1:200 Alexa Fluor 700-conjugated streptavidin (Invitrogen S21383) to detect bound ACE2. Cells were washed and resuspended in PBS-BSA for flow cytometry.

At each ACE2 sample concentration, single RBD^+^ cells were partitioned into bins of ACE2 binding (Alexa Fluor 700 fluorescence) as shown in Figure 1E using a BD FACSAria II. A minimum of 6.5 million cells were collected at each sample concentration, and sorted into SD -Ura -Trp with pen-strep antibiotic and 1% BSA. Collected cells in each bin were grown overnight in 1 mL SD -Ura -Trp + pen-strep, and plasmid was isolated using a 96-well yeast miniprep kit (Zymo D2005) according to kit instructions, with the addition of an extended (>2 hr) Zymolyase treatment and a -80°C freeze/thaw prior to cell lysis. N16 barcodes in each post-sort sample were PCR amplified as described in [4] and submitted for Illumina sequencing. Barcode sequence reads are available on the NCBI SRA, BioProject PRJNA770094, BioSample SAMN56444866.

Demultiplexed Illumina barcode reads were matched to library barcodes in barcode-mutant lookup tables using dms_variants (version 1.4.3), yielding a table of counts of each barcode in each FACS bin, available at https://github.com/tstarrlab/SARS-CoV-2-RBD_DMS_Omicron-KP3-LP8/blob/main/results/counts/variant_counts.csv. Read counts in each FACS bin were downweighted by the ratio of total sequence reads from a bin to the number of cells that were sorted into that bin from the FACS log.

We estimated the level of ACE2 binding of each barcoded mutant at each ACE2 concentration based on its distribution of counts across FACS bins as the simple mean bin [4]. We determined the ACE2-binding constant *K*_D_ for each barcoded mutant via nonlinear least-squares regression using the standard non-cooperative Hill equation relating the mean sort bin to the ACE2 labeling concentration and free parameters *a* (titration response range) and *b* (titration curve baseline):

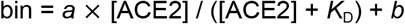

The measured mean bin value for a barcode at a given ACE2 concentration was excluded from curve fitting if fewer than 2 counts were observed across the four FACS bins or if counts exhibited bimodality (>40% of counts of a barcode were found in each of two non-consecutive bins). To avoid errant fits, we constrained the value *b* to (1, 2), *a* to (2, 3), and *K*_D_ to (10^-15^, 10^-5^). The fit for a barcoded variant was discarded if the average cell count across all sample concentrations was below 2, or if more than one sample concentration was missing. We also discarded curve fits where the normalized mean square residual (residuals normalized relative to the fit response parameter *a*) was >20 times the median value across all titration fits. Final binding constants were expressed as -log_10_(*K*_D_), where higher values indicate higher binding affinity. The complete computational pipeline for calculating and filtering per-barcode binding constants is available at https://github.com/tstarrlab/SARS-CoV-2-RBD_DMS_Omicron-KP3-LP8/blob/main/results/summary/compute_binding_Kd.md, and per-barcode affinity values are available at https://github.com/tstarrlab/SARS-CoV-2-RBD_DMS_Omicron-KP3-LP8/blob/main/results/binding_Kd/bc_binding.csv.

The affinity measurements of replicate barcodes representing an identical amino acid mutant were averaged within each experimental duplicate. The correlations in collapsed affinities in each duplicate experiment are shown in Figure 1G. The final measurement was determined as the average of duplicate measurements. The final -log_10_(*K*_D_) for each mutant and number of replicate barcode collapsed into this final measurement for each RBD mutant are given in Supplemental Data 1 and https://github.com/tstarrlab/SARS-CoV-2-RBD_DMS_Omicron-KP3-LP8/blob/main/results/final_variant_scores/final_variant_scores.csv, which includes data also from prior SARS-CoV-2 variant DMS datasets [33,36,41].

### RBD expression deep mutational scanning

Pooled libraries were grown and induced for RBD expression as described above. Induced cells were washed and labeled with 1:100 FITC-conjugated chicken anti-Myc to label for RBD expression via a C-terminal Myc tag, and washed in preparation for FACS. Single cells were partitioned into bins of RBD expression (FITC fluorescence) using a BD FACSAria II as shown in Figure 1F. A total of >10 million viable cells (estimated by plating dilutions of post-sort samples) were collected across bins for each library. Cells in each bin were grown out in SD -Ura -Trp + pen-strep + 1% BSA, plasmid isolated, and N16 barcodes sequenced as described above. Barcode reads are available on the NCBI SRA, BioProject PRJNA770094, BioSample SAMN56444866.

Demultiplexed Illumina barcode reads were matched to library barcodes in barcode-mutant lookup tables using dms_variants (version 1.4.3), yielding a table of counts of each barcode in each FACS bin, available at https://github.com/tstarrlab/SARS-CoV-2-RBD_DMS_Omicron-KP3-LP8/blob/main/results/counts/variant_counts.csv. Read counts in each bin were downweighted using the post-sort colony counts instead of the FACS log counts as with ACE2 titrations above to account for unequal viability of cells in FITC fluorescence bins (i.e., many cells in bin 1 are non-expressing because they have lost the low-copy expression plasmid and do not grow out post-FACS in selective media).

We estimated the level of RBD expression (log-mean fluorescence intensity, logMFI) of each barcoded mutant based on its distribution of counts across FACS bins and the known log-transformed fluorescence boundaries of each sort bin using a maximum likelihood approach [4,60] implemented via the fitdistrplus package in R [61]. Expression measurements were discarded for barcodes for which fewer than 10 counts were observed across the four FACS bins. The full pipeline for computing per-barcode expression values is available at https://github.com/tstarrlab/SARS-CoV-2-RBD_DMS_Omicron-KP3-LP8/blob/main/results/summary/compute_expression_meanF.md and per-barcode expression measurements are available at https://github.com/tstarrlab/SARS-CoV-2-RBD_DMS_Omicron-KP3-LP8/blob/main/results/expression_meanF/bc_expression.csv. Final mutant expression values were collapsed within and across replicates as described above, with correlation between experimental replicates shown in Figure 1H. Final mutant expression values and number of replicate barcode collapsed into this final measurement for each RBD mutant are available in Supplemental Data 1 and at https://github.com/tstarrlab/SARS-CoV-2-RBD_DMS_Omicron-KP3-LP8/blob/main/results/final_variant_scores/final_variant_scores.csv.

### Quantification of epistasis

Epistatic shifts at each site between pairs of RBD variants were quantified as described by [33]. Briefly, affinity phenotypes of each mutant at a site were transformed to a probability analog via a Boltzmann weighting, and the “epistatic shift” metric was calculated as the Jensen-Shannon divergence between the vectors of 21 amino acid probabilities (including the deletion character, when present). The Jensen-Shannon divergence ranges from 0 for two vectors of probabilities that are identical to 1 for two vectors that are completely dissimilar. To avoid noisier measurements artifactually inflating the epistatic shift metric, a given amino acid mutation was only included in the computation if it was sampled with a minimum of 3 replicate barcodes in each RBD background being compared. The calculation of epistatic shifts can be found at https://github.com/tstarrlab/SARS-CoV-2-RBD_DMS_Omicron-KP3-LP8/blob/main/results/summary/epistatic_shifts.md.

## Acknowledgements

We thank Bernadeta Dadonaite and Jesse Bloom for helpful discussion. We thank the Flow Cytometry Core Facility at the University of Utah Health Sciences Campus supported by NIH (5P30CA042014-24), and the University of Utah Center for High Performance Computing supported by NIH (1S10OD021644-01A1).

## Data and Materials Availability

Site saturation mutagenesis libraries and respective isogenic parental plasmid stocks are available from Addgene (Addgene ID in process). Raw sequencing data are available from the NCBI SRA under BioProject PRJNA770094, BioSamples SAMN56444854 (PacBio sequencing) and SAMN56444866 (Illumina barcode sequencing). All code and data at various stages of processing is available at: https://github.com/tstarrlab/SARS-CoV-2-RBD_DMS_Omicron-KP3-LP8/tree/main. Final mutant deep mutational scanning measurements are available in Supplemental Data 1.

## REFERENCES

1. Carabelli AM, Peacock TP, Thorne LG, Harvey WT, Hughes J, COVID-19 Genomics UK Consortium, et al. SARS-CoV-2 variant biology: immune escape, transmission and fitness. Nat Rev Microbiol. 2023;21: 162–177.

2. Piccoli L, Park Y-J, Tortorici MA, Czudnochowski N, Walls AC, Beltramello M, et al. Mapping Neutralizing and Immunodominant Sites on the SARS-CoV-2 Spike Receptor-Binding Domain by Structure-Guided High-Resolution Serology. Cell. 2020;183: 1024–1042.e21.

3. Starr TN, Czudnochowski N, Liu Z, Zatta F, Park Y-J, Addetia A, et al. SARS-CoV-2 RBD antibodies that maximize breadth and resistance to escape. Nature. 2021;597: 97–102.

4. Starr TN, Greaney AJ, Hilton SK, Ellis D, Crawford KHD, Dingens AS, et al. Deep Mutational Scanning of SARS-CoV-2 Receptor Binding Domain Reveals Constraints on Folding and ACE2 Binding. Cell. 2020;182: 1295–1310.e20.

5. Lynch RM, Wong P, Tran L, O’Dell S, Nason MC, Li Y, et al. HIV-1 fitness cost associated with escape from the VRC01 class of CD4 binding site neutralizing antibodies. J Virol. 2015;89: 4201–4213.

6. Wrapp D, Wang N, Corbett KS, Goldsmith JA, Hsieh C-L, Abiona O, et al. Cryo-EM structure of the 2019-nCoV spike in the prefusion conformation. Science. 2020;367: 1260–1263.

7. Walls AC, Park Y-J, Tortorici MA, Wall A, McGuire AT, Veesler D. Structure, Function, and Antigenicity of the SARS-CoV-2 Spike Glycoprotein. Cell. 2020;181: 281–292.e6.

8. Tse AL, Acreman CM, Ricardo-Lax I, Berrigan J, Lasso G, Balogun T, et al. Distinct pathways for evolution of enhanced receptor binding and cell entry in SARS-like bat coronaviruses. PLoS Pathog. 2024;20: e1012704.

9. Tse AL, Lasso G, Berrigan J, McLellan JS, Chandran K, Miller EH. Bat sarbecovirus WIV1-CoV bears an adaptive mutation that alters spike dynamics and enhances ACE2 binding. PLoS Pathog. 2025;21: e1013123.

10. Wang J, Ma Y, Li Z, Yuan H, Liu B, Li Z, et al. SARS-related coronavirus S-protein structures reveal synergistic RBM interactions underpinning high-affinity human ACE2 binding. Sci Adv. 2025;11: eadr8772.

11. Kang L, He G, Sharp AK, Wang X, Brown AM, Michalak P, et al. A selective sweep in the Spike gene has driven SARS-CoV-2 human adaptation. Cell. 2021;184: 4392–4400.e4.

12. Zhang S, Liang Q, He X, Zhao C, Ren W, Yang Z, et al. Loss of Spike N370 glycosylation as an important evolutionary event for the enhanced infectivity of SARS-CoV-2. Cell Res. 2022;32: 315–318.

13. Gobeil SM-C, Henderson R, Stalls V, Janowska K, Huang X, May A, et al. Structural diversity of the SARS-CoV-2 Omicron spike. Mol Cell. 2022;82: 2050–2068.e6.

14. Zhang QE, Lindenberger J, Parsons RJ, Thakur B, Parks R, Park CS, et al. SARS-CoV-2 Omicron XBB lineage spike structures, conformations, antigenicity, and receptor recognition. Mol Cell. 2024;84: 2747–2764.e7.

15. Korber B, Fischer W, Theiler J. Real-time monitoring of SARS-CoV-2 evolution during the COVID-19 pandemic. Cell Host Microbe. 2025;33: 1802–1806.

16. DeGrace MM, Ghedin E, Frieman MB, Krammer F, Grifoni A, Alisoltani A, et al. Defining the risk of SARS-CoV-2 variants on immune protection. Nature. 2022;605: 640–652.

17. Fowler DM, Fields S. Deep mutational scanning: a new style of protein science. Nat Methods. 2014;11: 801–807.

18. Greaney AJ, Starr TN, Gilchuk P, Zost SJ, Binshtein E, Loes AN, et al. Complete Mapping of Mutations to the SARS-CoV-2 Spike Receptor-Binding Domain that Escape Antibody Recognition. Cell Host Microbe. 2021;29: 44–57.e9.

19. Ouyang WO, Tan TJC, Lei R, Song G, Kieffer C, Andrabi R, et al. Probing the biophysical constraints of SARS-CoV-2 spike N-terminal domain using deep mutational scanning. Sci Adv. 2022;8: eadd7221.

20. Taft JM, Weber CR, Gao B, Ehling RA, Han J, Frei L, et al. Deep mutational learning predicts ACE2 binding and antibody escape to combinatorial mutations in the SARS-CoV-2 receptor-binding domain. Cell. 2022;185: 4008–4022.e14.

21. Kugathasan R, Sukhova K, Moshe M, Kellam P, Barclay W. Deep mutagenesis scanning using whole trimeric SARS-CoV-2 spike highlights the importance of NTD-RBD interactions in determining spike phenotype. PLoS Pathog. 2023;19: e1011545.

22. Francino-Urdaniz IM, Steiner PJ, Kirby MB, Zhao F, Haas CM, Barman S, et al. One-shot identification of SARS-CoV-2 S RBD escape mutants using yeast screening. Cell Rep. 2021;36: 109627.

23. Dadonaite B, Crawford KHD, Radford CE, Farrell AG, Yu TC, Hannon WW, et al. A pseudovirus system enables deep mutational scanning of the full SARS-CoV-2 spike. Cell. 2023;186: 1263–1278.e20.

24. Cao Y, Wang J, Jian F, Xiao T, Song W, Yisimayi A, et al. Omicron escapes the majority of existing SARS-CoV-2 neutralizing antibodies. Nature. 2022;602: 657–663.

25. Starr TN, Greaney AJ, Addetia A, Hannon WW, Choudhary MC, Dingens AS, et al. Prospective mapping of viral mutations that escape antibodies used to treat COVID-19. Science. 2021;371: 850–854.

26. Rosen LE, Tortorici MA, De Marco A, Pinto D, Foreman WB, Taylor AL, et al. A potent pan-sarbecovirus neutralizing antibody resilient to epitope diversification. Cell. 2024;187: 7196–7213.e26.

27. Thadani NN, Gurev S, Notin P, Youssef N, Rollins NJ, Ritter D, et al. Learning from prepandemic data to forecast viral escape. Nature. 2023;622: 818–825.

28. Maher MC, Bartha I, Weaver S, di Iulio J, Ferri E, Soriaga L, et al. Predicting the mutational drivers of future SARS-CoV-2 variants of concern. Sci Transl Med. 2022;14: eabk3445.

29. Lamb KD, Hughes J, Lytras S, Young F, Koci O, Herzig JC, et al. From single-sequences to evolutionary trajectories: protein language models capture the evolutionary potential of SARS-CoV-2. Nat Commun. 2026;1–16.

30. Hie B, Zhong ED, Berger B, Bryson B. Learning the language of viral evolution and escape. Science. 2021;371: 284–288.

31. Starr TN, Thornton JW. Epistasis in protein evolution. Protein Sci. 2016;25: 1204–1218.

32. Zahradník J, Marciano S, Shemesh M, Zoler E, Harari D, Chiaravalli J, et al. SARS-CoV-2 variant prediction and antiviral drug design are enabled by RBD in vitro evolution. Nat Microbiol. 2021;6: 1188–1198.

33. Starr TN, Greaney AJ, Hannon WW, Loes AN, Hauser K, Dillen JR, et al. Shifting mutational constraints in the SARS-CoV-2 receptor-binding domain during viral evolution. Science. 2022;377: 420–424.

34. Moulana A, Dupic T, Phillips AM, Chang J, Nieves S, Roffler AA, et al. Compensatory epistasis maintains ACE2 affinity in SARS-CoV-2 Omicron BA.1. Nat Commun. 2022;13: 7011.

35. Jian F, Feng L, Yang S, Yu Y, Wang L, Song W, et al. Convergent evolution of SARS-CoV-2 XBB lineages on receptor-binding domain 455-456 synergistically enhances antibody evasion and ACE2 binding. PLoS Pathog. 2023;19: e1011868.

36. Taylor AL, Starr TN. Deep mutational scans of XBB.1.5 and BQ.1.1 reveal ongoing epistatic drift during SARS-CoV-2 evolution. PLoS Pathog. 2023;19: e1011901.

37. Mellis IA, Wu M, Hong H, Tzang C-C, Bowen A, Wang Q, et al. Antibody evasion and receptor binding of SARS-CoV-2 LP.8.1.1, NB.1.8.1, XFG, and related subvariants. Cell Rep. 2025;44: 116440.

38. Kaku Y, Uriu K, Okumura K, Genotype to Phenotype Japan (G2P-Japan) Consortium, Ito J, Sato K. Virological characteristics of the SARS-CoV-2 KP.3.1.1 variant. Lancet Infect Dis. 2024;24: e609.

39. Chen L, Kaku Y, Okumura K, Uriu K, Zhu Y, Genotype to Phenotype Japan (G2P-Japan) Consortium, et al. Virological characteristics of the SARS-CoV-2 LP.8.1 variant. Lancet Infect Dis. 2025;25: e193.

40. Adams RM, Mora T, Walczak AM, Kinney JB. Measuring the sequence-affinity landscape of antibodies with massively parallel titration curves. Elife. 2016;5. doi:10.7554/eLife.23156

41. Starr TN, Greaney AJ, Stewart CM, Walls AC, Hannon WW, Veesler D, et al. Deep mutational scans for ACE2 binding, RBD expression, and antibody escape in the SARS-CoV-2 Omicron BA.1 and BA.2 receptor-binding domains. PLoS Pathog. 2022;18: e1010951.

42. Taylor AL, Starr TN. Deep mutational scanning of SARS-CoV-2 Omicron BA.2.86 and epistatic emergence of the KP.3 variant. Virus Evol. 2024;10: veae067.

43. Pollock DD, Thiltgen G, Goldstein RA. Amino acid coevolution induces an evolutionary Stokes shift. Proc Natl Acad Sci U S A. 2012;109: E1352–9.

44. Shah P, McCandlish DM, Plotkin JB. Contingency and entrenchment in protein evolution under purifying selection. Proc Natl Acad Sci U S A. 2015;112: E3226–35.

45. Dadonaite B, Harari S, Larsen BB, Kampman L, Harteloo A, Elias-Warren A, et al. Spike mutations that affect the function and antigenicity of recent KP.3.1.1-like SARS-CoV-2 variants. Liu S-L, editor. J Virol. 2025;99. doi:10.1128/jvi.01423-25

46. Liu C, Zhou D, Dijokaite-Guraliuc A, Supasa P, Duyvesteyn HME, Ginn HM, et al. A structure-function analysis shows SARS-CoV-2 BA.2.86 balances antibody escape and ACE2 affinity. Cell Rep Med. 2024;5: 101553.

47. Wang Q, Iketani S, Li Z, Guo Y, Yeh AY, Liu M, et al. Antigenic characterization of the SARS-CoV-2 Omicron subvariant BA.2.75. Cell Host & Microbe. 2022;30: 1512–1517.e4.

48. Cao Y, Jian F, Wang J, Yu Y, Song W, Yisimayi A, et al. Imprinted SARS-CoV-2 humoral immunity induces convergent Omicron RBD evolution. Nature. 2023;614: 521–529.

49. Harbison AM, Fogarty CA, Phung TK, Satheesan A, Schulz BL, Fadda E. Fine-tuning the spike: role of the nature and topology of the glycan shield in the structure and dynamics of the SARS-CoV-2 S. Chem Sci. 2022;13: 386–395.

50. Cerutti G, Guo Y, Liu L, Liu L, Zhang Z, Luo Y, et al. Cryo-EM structure of the SARS-CoV-2 Omicron spike. Cell Reports. 2022;38: 110428.

51. Wieczór M, Tang PK, Orozco M, Cossio P. Omicron mutations increase interdomain interactions and reduce epitope exposure in the SARS-CoV-2 spike. iScience. 2023;26: 105981.

52. Feng Z, Huang J, Baboo S, Diedrich JK, Bangaru S, Paulson JC, et al. Structural and functional insights into the evolution of SARS-CoV-2 KP.3.1.1 spike protein. Cell Rep. 2025;44: 115941.

53. Cao Y, Jian F, Zhang Z, Yisimayi A, Hao X, Bao L, et al. Rational identification of potent and broad sarbecovirus-neutralizing antibody cocktails from SARS convalescents. Cell Rep. 2022;41: 111845.

54. Poelwijk FJ, Socolich M, Ranganathan R. Learning the pattern of epistasis linking genotype and phenotype in a protein. Nat Commun. 2019;10: 4213.

55. Park Y, Metzger BPH, Thornton JW. The simplicity of protein sequence-function relationships. Nat Commun. 2024;15: 7953.

56. Greaney AJ, Starr TN, Eguia RT, Loes AN, Khan K, Karim F, et al. A SARS-CoV-2 variant elicits an antibody response with a shifted immunodominance hierarchy. PLoS Pathog. 2022;18: e1010248.

57. Wentz AE, Shusta EV. A novel high-throughput screen reveals yeast genes that increase secretion of heterologous proteins. Appl Environ Microbiol. 2007;73: 1189–1198.

58. Gietz RD, Schiestl RH. Large-scale high-efficiency yeast transformation using the LiAc/SS carrier DNA/PEG method. Nat Protoc. 2007;2: 38–41.

59. Crawford KHD, Bloom JD. alignparse: A Python package for parsing complex features from high-throughput long-read sequencing. J Open Source Softw. 2019;4. doi:10.21105/joss.01915

60. Peterman N, Levine E. Sort-seq under the hood: implications of design choices on large-scale characterization of sequence-function relations. BMC Genomics. 2016;17: 206.

61. Delignette-Muller M, Dutang C. fitdistrplus: An R Package for Fitting Distributions. Journal of Statistical Software, Articles. 2015;64: 1–34.

